# Repurposing of Miglustat to inhibit the coronavirus Severe Acquired Respiratory Syndrome SARS-CoV-2

**DOI:** 10.1101/2020.05.18.101691

**Authors:** Sreejith Rajasekharan, Rafaela Milan Bonotto, Yvette Kazungu, Lais Nascimento Alves, Monica Poggianella, Pamela Martinez Orellana, Natasa Skoko, Sulena Polez, Alessandro Marcello

**Affiliations:** Laboratory of Molecular Virology, International Centre for Genetic Engineering and Biotechnology (ICGEB) Padriciano, 99 - 34149 Trieste, ITALY; Biotechnology Development, International Centre for Genetic Engineering and Biotechnology (ICGEB) Padriciano, 99 - 34149 Trieste, ITALY

**Author notes:** These authors contributed equally to this work (Sreejith Rajasekharan, Rafaela Milan Bonotto).

**Keywords:** COVID-19, SARS-CoV-2, coronavirus, inhibitor, antiviral, Miglustat, iminosugar, Zavesca, NB-DNJ, Spike

## Abstract

Repurposing clinically available drugs to treat the new coronavirus disease COVID-19 is an urgent need in these early stages of the SARS-CoV-2 pandemic, when very few treatment options are available. The iminosugar Miglustat is a well-characterized drug for the treatment of rare genetic lysosome storage diseases such as Gaucher and Niemann-Pick type C, and has also been described to be active against a variety of enveloped viruses. The activity of Miglustat is here demonstrated for SARS-CoV-2 at concentrations achievable in the plasma by current clinical regimens without cytotoxicity. The drug acts at the post-entry level and leads to a marked decrease of viral proteins and release of infectious virus. The mechanism resides in the inhibitory activity towards α-glucosidases that are involved in early stages of glycoprotein N-linked oligosaccharide processing in the endoplasmic reticulum, leading to a marked decrease of the viral Spike protein. The wealth of available data on the clinical use of Miglustat for the treatment of lysosomal storage disorders and the antiviral properties against SARS-CoV-2 make it an ideal candidate for drug repurposing.

## Introduction

The novel Severe Acquired Respiratory Syndrome (SARS-CoV-2) coronavirus, the etiologic agent of coronavirus disease 2019 (COVID-19), has now spread in everywhere causing a global pandemic (1, 2). To date there have been more than 3.5 million reported cases and 250.000 deaths worldwide urging for a global effort to tackle the disease (3). SARS-CoV-2 belongs to the genus *Betacoronavirus* of the order/family/sub-family *Nidovirales*/*Coronaviridae/Coronavirinae*. The virion is enveloped and contains a single RNA genome of positive polarity. Morphologically, SARS-CoV-2 is about 120 nm in diameter with large projections of the heavily glycosylated trimeric spike (S) proteins. Other surface proteins include the membrane (M) and envelope (E) proteins, while inside the envelope the helical nucleocapsid (N) wraps the viral RNA. The virus targets cells of the upper and lower respiratory tract epithelia through the viral Spike that binds to the angiotensin-converting enzyme 2 (ACE2) receptor, a process facilitated by the host type 2 transmembrane serine protease, TMPRSS2. Once inside the cell, viral polyproteins are synthesized that encode for the replication machinery required to synthesize new RNA via its RNA-dependent RNA polymerase activity. Replication is cytoplasmatic at the level of the endoplasmic reticulum (ER) that is heavily rearranged. Structural proteins are then synthesized leading to completion of assembly and release of viral particles (4, 5). Currently, no specific treatment against SARS-CoV-2 is available and the only antiviral therapy comes from repurposing of drugs developed for other viral infections. Lopinavir and ritonavir, remdesivir, (hydroxy)chloroquine, umifenovir and favipiravir are examples that are currently being evaluated, but none have been conclusively shown to be effective (6).

The iminosugar Miglustat (Zavesca; N-butyl-1-deoxynojirimycin, NB-DNJ) inhibits α-glucosidases I and II that are involved in early stages of glycoprotein N-linked oligosaccharide processing in the ER (7). Because most enveloped viruses require glycosylation for surface protein folding and secretion, modulation of the oligosaccharide to induce a reduction in infectivity became a strategy for treatment of the immune deficiency virus type 1 (HIV-1), culminating in phase I/II clinical trials (8, 9). The use of iminosugars to misfold viral glycoprotein as a therapeutic approach has so far been applied to several other viral infections including: hepatitis B and C viruses, Dengue and other flaviviruses, and Ebola virus (10-12). An additional property of certain iminosugars is the Glucosyltransferase inhibition activity, which is the basis for current therapy of rare genetic lysosome storage diseases such as Gaucher and Niemann-Pick type C (13). This activity of Miglustat could impact virus entry by modification of the plasma membrane.

The wealth of data on the clinical use of Miglustat for the treatment of lysosomal storage disorders and the antiviral properties observed on enveloped viruses make Miglustat an ideal candidate of drug repurposing for COVID-19. In this work Miglustat is shown to be active for the inhibition of SARS-CoV-2 in different cell types at concentrations compatible with those obtained for the treatment of Gaucher and Niemann-Pick type C in patients. Time of addition studies indicate that the inhibitory activity is at the post-entry level and affects release of infectious virus. The proper folding and release of the Spike protein, and progeny virus appear mostly affected.

## Materials and Methods

### Cells, virus and antiviral assay

Vero E6 cells (ATCC-1586) HEK 293T (ATCC CRL-3216), A549 (ATCC CCL-185), U2OS (ATCC HTB-96) and human hepatocarcinoma Huh7 cells kindly provided by Ralf Bartenschlager (University of Heidelberg, Germany) were cultured in Dulbecco’s modified Eagle’s medium (DMEM) supplemented with 10% fetal bovine serum (FBS, Gibco) and antibiotics. Cell cultures were maintained at 37 °C under 5% CO_2_. Cells were routinely tested for mycoplasma contamination.

Working stocks of SARS-CoV-2 ICGEB-FVG_5 isolated in Trieste, Italy, were routinely propagated and titrated on Vero E6 cells (14). Plaque assay was performed by incubating dilutions of SARS-CoV-2 on Vero E6 monolayers at 37 °C for 1 hour, which were then washed with phosphate buffered saline (PBS) and overlaid with DMEM 2% FBS containing 1.5% carboxymethylcellulose for 3 days. Cells were then fixed with 3.7% paraformaldehyde (PFA) and stained with crystal violet 1%. Cytotoxicity assay was performed with Alamar Blue (ThermoFisher) according to manufacturer’s instructions.

### Drugs and proteins

Miglustat (NB-DNJ) was purchased from Sigma (B8299). The drug was dissolved in DMSO to obtain a stock solution, while following dilutions were made directly in growth medium.

The SARS-CoV-2 Spike protein receptor-binding domain (RBD) was expressed from pCAGGS using a construct generously provided by Florian Krammer (Mount Sinai, New York) (15). Plasmid was transfected in 293T cells and cell extracts and supernatants were harvested 24 hours post-transfection. Miglustat 200 µM was added after transfection and maintained in the medium until the end of the experiment.

Sequence coding for the full length Spike protein was obtained from isolate Wuhan-Hu-1 (NCBI Reference Sequence: NC_045512.2). The nucleotide sequence, fused to an immunoglobulin leader sequence (sec) at the N-terminus, codon optimized for expression in mammalian cells, was obtained as synthetic DNA fragment from GenScript Biotech (Netherlands) and cloned as HindIII/ApaI into a pCDNA3 vector.

### Plaque reduction assay

Vero E6 cells were seeded at 6×10^4^ cells/well density in a 48 wells’ plate and incubated at 37 °C overnight. Cells were infected with 30 viral PFU/well, and incubated at 37 °C for 1 hour. Following incubation, the virus was removed and the wells washed with 1x PBS. The infected cells were maintained with 800 µL of overlay medium containing 1.5% carboxymethylcellulose (CMC) with DMEM + 2% heat-inactivated FBS, and Miglustat dilutions. Cells were then incubated at 37 °C for 3 days. Finally, cells were fixed with 3.7% PFA and stained with crystal violet. Plaques were counted and values were normalized to vehicle (DMSO). The plaque reduction assays were conducted in double replicates for three independent experiments. Inhibition was calculated with the formula: (1-(average plaques Miglustat/average plaques Vehicle))*100 and plotted against dilutions as antilog. For the cytotoxicity assay, fluorescence readings were normalized for vehicle and percent plotted against dilutions. The half maximal effective concentration (EC_50_) and cytotoxic concentration (CC_50_) were calculated using GraphPad Prism Version 7.

### Immunoblotting and Immunofluorescence

A recombinant monoclonal reactive with the receptor-binding domain of the S protein was generated based on a mouse small immune protein (SIP) scaffold (mSIP-3022). The DNA fragment encoding for the variable regions VL (NCBI accession code: DQ168570.1) and VH (NCBI accession code: DQ168569.1) of human monoclonal antibody, clone CR3022 (16, 17), was synthetized as scFV by GenScript Biotech (Netherlands) and cloned into ApaLI-BspEI sites upstream of Hinge-CH2-CH3 domains of mouse IgG2b expression vector as described previously (18). The plasmid was transfected in ExpiCHO-S cells (Gibco-Thermo Fisher Scientific) following manufacturer’s instructions. Eight days post transfection the supernatant was loaded on the HiTrap protein G HP 1ml column (GE Healthcare) in binding buffer (20 mM sodium phosphate pH 7.0) and eluted with acetic acid 50mM pH 2.7. Eluted antibody was immediately neutralized with 1M Tris pH 8, and analyzed by RP-HPLC and by SDS PAGE to maintain the dimeric structure. Production yield was 0.8 mg/ml. For immunofluorescence cells were fixed in 3.7% PFA, permeabilized with 0.01% Triton and processed with mSIP-3022 as per standard procedure (19). Since mSIP-3022 did not react with the denatured S protein a convalescent serum from a COVID-19 patient was used for immunoblotting at a 1:200 dilution. Images were acquired on a Zeiss LSM880 confocal microscope. For immunoblotting, whole-cell lysates were resolved by 12% SDS-PAGE and blotted onto nitrocellulose membranes. The membranes were blocked in 5% nonfat milk in Tris-buffered saline (TBS) plus 0.1% Tween 20 (TBST), followed by incubation with the human serum diluted 1:200 in the same solution for 1 hour at room temperature. After washing three times with TBST, secondary horseradish peroxidase (HRP)-conjugated antibodies were incubated for 1 hour at room temperature. The blots were developed using a chemioluminescent HRP substrate (Millipore). The anti-his antibody (monoclonal #8722 Sigma) was used at 1/2000 concentration for immunoblot.

### Viral RNA quantification

For real-time quantitative reverse transcription PCR (RT-qPCR) total cellular RNA was extracted with Trizol (Sigma) or QiAmp (Qiagen) according to the manufacturer’s protocol and treated with DNase I (Invitrogen). 500 ng were then reverse-transcribed with random primers and MMLV Reverse Transcriptase (Invitrogen). Quantification of mRNA was obtained by real-time PCR using the Kapa Sybr fast qPCR kit on a CFX96 Bio-Rad thermocycler. Primers for amplification were those listed by China CDC as ORF1ab and genomes were calculated on a standard quantified RNA (20).

### Statistics

Typically three independent experiments in triplicate repeats were conducted for each condition examined. Average values are shown with standard deviation and p-values, measured with a paired two-tailed t-test. Only significant pvalues are indicated by the asterisks above the graphs (**p < 0.01 highly significant; *p < 0.05 significant). Where asterisks are missing the differences are calculated as non-significant (n.s).

## Results

The antiviral properties of Miglustat were assessed by performing a plaque assay. SARS-CoV-2 strain ICGEB-FVG_5 was used to infect Vero E6 for 1 hour. After removal of the inoculum and wash in PBS, the cells were overlaid with medium containing 1.5% CMC and dilutions of drug as indicated in Figure 1A. 72 hours post-infection cells were fixed and stained to reveal plaques, which were visually counted. Data are from three independent experiments each in duplicate biological replicates. In parallel, cytotoxicity was assessed by the Alamar blue method at the indicated dilutions of drug. The effective concentration for 50% inhibition (EC_50_) of Miglustat was 41±22 µM with no apparent cyototoxicity until 1 mM (CC_50_ > 1 mM). The plaque assay was performed in Vero E6 cells, which are standard for the growth of SARS-CoV-2. However, further analysis would be better performed in cells of human origin.

**Figure 1.**
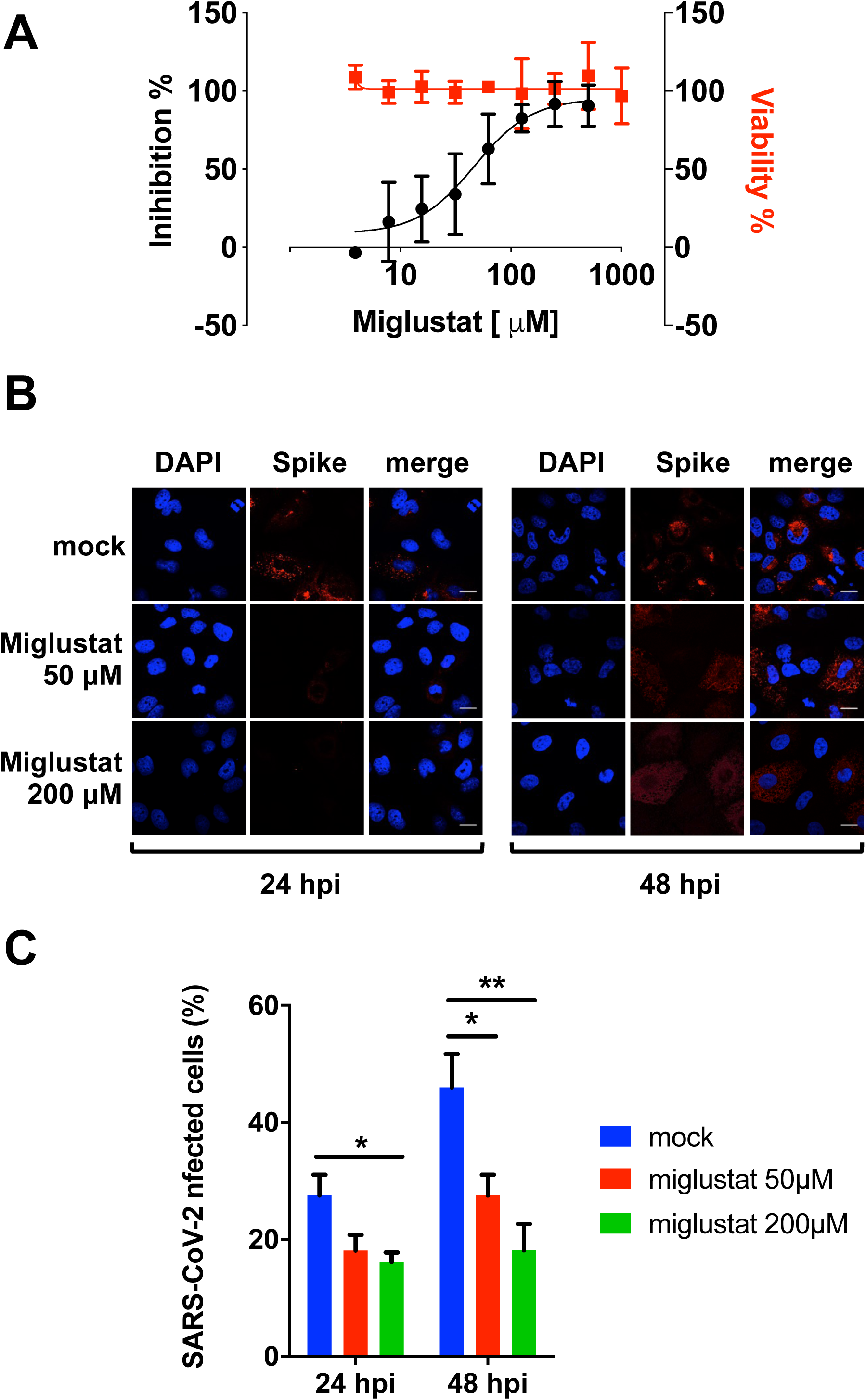
A) Antiviral plaque assay. Miglustat at the indicated concentrations was added to Vero E6 monolayers infected with SARS-CoV-2. Following incubation for three days cells were fixed and stained to count viral plaques against vehicle control, which were plotted as percent inhibitory activity (black dots). Cytotoxicity was measured by the Alamar blue method and data plotted as percent viability (red squares). B) Immunofluorescence assay. Huh7 cells were infected with SARS-CoV-2 moi = 0.1 and incubated with Miglustat as indicated. Cells were then fixed and stained with mSIP-3022 antibody against Spike (red) to acquire confocal images. Nuclei are stained by DAPI. Bar corresponds to 20 µm. C) Quantification of infected cells. SARS-CoV-2 infected cells were counted. Results from 200 cells per condition were plotted as percent infected cells.

In order to establish an infectious model in human cells a number of available cell lines were tested including U2OS (osteosarcoma), A549 (adenocarcinoma of the human alveolar basal epithelial), HEK 293 (human embryonic kidney cells) and Huh7 (hepatocellular carcinoma). None of the cell lines tested supported SARS-CoV-2 infection except for the Huh7 hepatocellular carcinoma cell line. As shown in Supplementary Figure S1A, Huh7 supported SARS-CoV-2 infection, albeit delayed 24 hours and at a lower efficiency compared to Vero E6. Furthermore, in order to stain infected cells a protocol for immunofluorescence was established. To this end, a recombinant monoclonal based on SIP mouse scaffold (mSIP-3022) was generated carrying the CDR regions of the antibody CR3022 reactive against the receptor binding domain (RBD) of the Spike protein of SARS-CoV-1, which showed high binding affinity also for SARS-CoV-2 Spike protein (17). As shown in Supplementary Figure S1B, mSIP-3022 efficiently stains the Spike protein in the cytoplasm of infected Vero E6 and Huh7 cells 24 hours post-infection.

Next, the efficiency of Huh7 infection was assessed in the presence of Miglustat. As shown in Figure 1B and quantified in Figure 1C, Miglustat maintained the number of Huh7 infected cells at the level observed 24 hpi, while mock treated cells showed an increase of infected cells at 48 hpi as expected form an expansion of the infection in the cell culture. These data confirm the inhibitory effect of Miglustat in cells of human origin and might suggest that the activity of Miglustat is at the level of the secretion of new infectious virus and not at the entry/replication level.

To better characterize this hypothesis, a time-of-addition (TOA) experiment was performed. Different conditions were used: pre-treatment, co-treatment and post-treatment. Huh7 cells were pre-treated with 200 µM Miglustat for 3h and then infected for 1h in the absence of drug (moi = 0.1). Afterwards, the virus was removed and the cells were cultured in drug-free medium until the end of the experiment. For co-treatment, the drug was added together with the virus during infection and then cells were maintained in drug-free medium. For post-entry experiment, drug was added at 3 h post-infection and maintained until the end of the experiment. As shown in Figure 2A, drug didn’t affect viral entry, neither was virucidal when administered concomitant with infection. Replication (intracellular viral RNA) was slightly affected at 48hpi and significantly at 72hpi consistent with the idea that Miglustat was effective at the post-entry level. This was reflected by the reduction of intracellular nucleocapsid N protein observed at both time points (Figure 2B). Interestingly, a strong reduction of infectious virus was observed in post-entry conditions (Figure 2C and 2D) paralleled by a decrease of extracellular viral genomes (Figure 2E and 2F) and N protein (Figure 2G).

**Figure 2.**
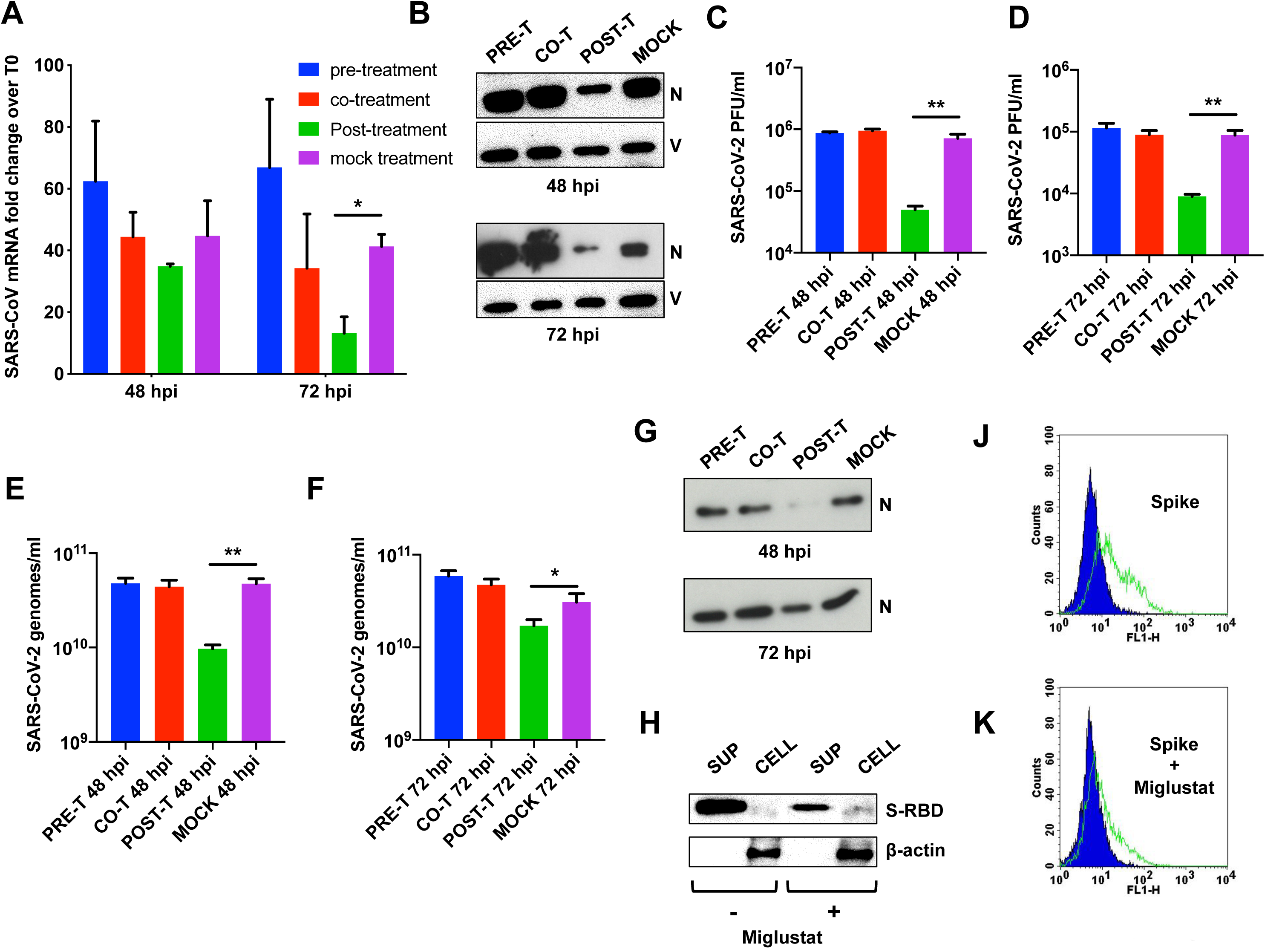
A) Time-of-addition experiment: SARS-CoV-2 genomic RNA. Huh7 cells were infected at moi = 0.1 and incubated with Miglustat before infection (pre-treatment), during infection (co-treatment) and after infection (post-treatment) as described in the text. At the indicated time points, total RNA was extracted from the infected cells and analyzed by RT PCR. Data are shown as fold-change normalized to their respective T0 values. B) Time-of-addition experiment: SARS-CoV-2 proteins. Protein extracts from Huh7 cells treated as in (A) were immunoblotted with a COVID-19 convalescent human serum. The N protein is indicated, with Vimentin as loading control. C-D) Time-of-addition experiment: SARS-CoV-2 infectious virus. The infectious virus produced in the experiment was measured as PFU/ml on Vero E6 cells as indicated. E-F) Time-of-addition experiment: SARS-CoV-2 secreted genomes. The SARS-CoV-2 genomes in the supernatant of infected cells were quantified by RT PCR as indicated. G) Time-of-addition experiment: SARS-CoV-2 secreted virions. The virion protein N of secreted SARS-CoV-2 was detected with a convalescent human serum. H) Secretion of SARS-CoV-2 Spike RBD. The his-tagged Spike-RBD was expressed in HEK 293T cells in the presence of Miglustat and protein detected by immunoblot for the his-tag both in supernatant and cell extracts with β-actin as loading control. J-K) Surface expression of SARS-CoV-2 Spike. Full-length SARS-CoV-2 Spike was expressed in HEK-293T cells in the presence of Miglustat and its expression and correct folding on the cell surface was detected with the mSIP-3022 antibody.

Miglustat activity at the post-entry level is expected to target proper folding and glycosylation of virion proteins. Spike and its receptor-binding domain are heavily glycosylated and undergo folding and glycosylation through the ER before being secreted and exposed on the plasma membrane. Previous data on the Spike protein of SARS-CoV-1 indicated that both glycosylation and secretion were affected by Miglustat (21, 22). Therefore we took advantage of an expression vector for SARS-CoV-2 Spike RBD to assess the effect of Miglustat treatment on protein release from transfected 293T cells. As shown in figure 2H, the protein is highly abundant in the cell supernatant in normal conditions, but, upon treatment with 200 µM Miglustat, the protein was much less abundant. These data point to a role of Miglustat as an inhibitor of the proper folding and release of functional Spike protein. To reinforce this observation, full-length spike was transfected in 293T cells and the fully folded protein expressed on the cell surface detected by the conformation-dependent mSIP-3022 antibody. As shown in Figure 2J and 2K, Miglustat reduce the amount of protein on the cell surface.

## Discussion

Host directed antiviral therapy is a strategy of inhibiting virus infection by targeting host factors that are essential for viral replication (23). Currently there is a pressing need for antiviral drugs for immediate use in the context of SARS-CoV-2 infection. Miglustat is a drug that is in current clinical use for the treatment of certain genetic disorders and has shown to be active against a variety of viral infections, making it a suitable candidate for drug repurposing towards SARS-CoV-2. In this work the activity of Miglustat against SARS-CoV-2 has been demonstrated in vitro with EC_50_ in the micromolar range. The standard dosage for lysosomal storage diseases such as Gaucher or Niemann Pick is 100 mg/3 times a day, with a maximum daily dose of 600 mg/day. A single dose of 100 mg Miglustat reaches a peak in plasma concentration of around 3-5 µM within 4 hours, while half-life is approximately 8 hours. If this dose is administered every 4 hours/6 times per day, the plasma concentration of Miglustat would stabilize around 10 µM (24). If 200 mg Miglustat is administered every 8 hours/3 times a day, the plasma concentration could be also higher than 10 µM in 24 hours. However, increased dosage could lead to well-described adverse reactions that include tremors, diarrhea, numbness and thrombocytopenia.

Miglustat has been shown to act through two different mechanisms: at the level of virus entry, by perturbing the plasma membrane, and at the level of folding and secretion of virion proteins, by affecting essential glycosylation steps in the ER. The latter mechanism is supported by the data presented in this work, where a strong antiviral activity is detected only when the drug is added post-infection. Correct folding and secretion of glycoproteins is a process tightly controlled in the ER by chaperons such as Calnexin that recognize specific glycosylation intermediates (25). Miglustat interferes with this process resulting in the accumulation of misfolded proteins and a defect in secretion. Consistently, the Spike protein of SARS-CoV-1 has been shown to bind Calnexin and disruption of this function caused decrease of virus infectivity (22).

In conclusion this work provides in vitro evidence for the use of Miglustat as inhibitor of SARS-CoV-2 and proposes its use in clinical trials for COVID-19 patients.

---

**Supplementary Figure S1.**
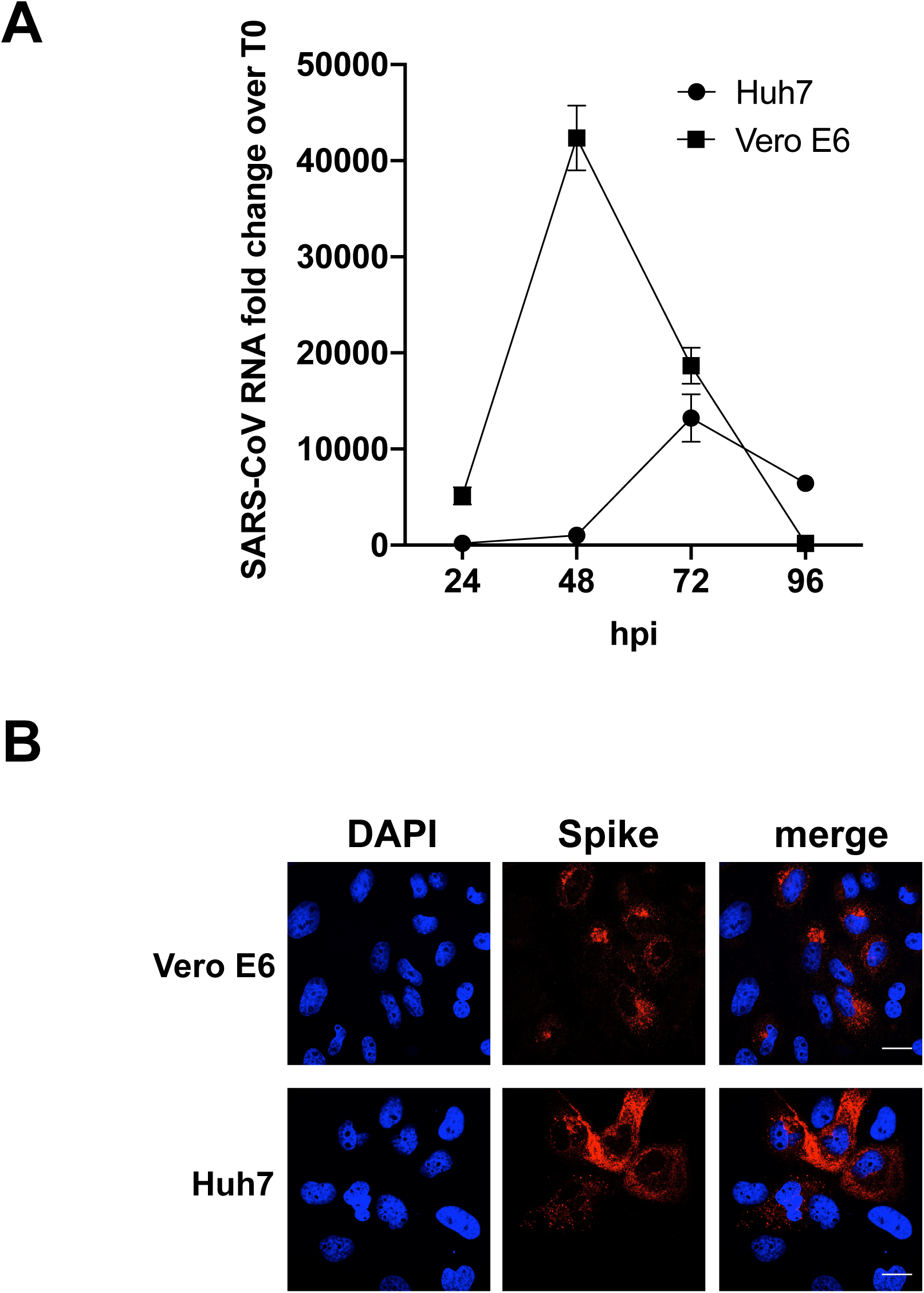
A) SARS-CoV-2 infectivity. Vero E6 and Huh7 cells were infected in parallel with SARS-CoV-2 at moi 0.1. At the indicated time points RNA was extracted from the cells and measured by RT PCR for SARS-CoV-2 genomes. Data are plotted as fold change over T0. B) Immunofluorecence assay. Vero E6 and Huh7 cells were infected with SARS-CoV-2 moi = 0.1 for 24 hours. Cells were then fixed and stained with mSIP-3022 antibody against Spike (red) to acquire confocal images. Nuclei are stained by DAPI. Bar corresponds to 20 µM.

## Acknowledgments

Work on SARS-CoV-2 in AM’s laboratory is supported by ICGEB intramural funds, by the Beneficientia Stiftung and by the SNAM Foundation. We thank Professor Bruno Bembi for helpful suggestions.

## Notes

### Competing Interest Statement

The authors have declared no competing interest.

## References

1. Zhou P, Yang XL, Wang XG, Hu B, Zhang L, Zhang W, Si HR, Zhu Y, Li B, Huang CL, Chen HD, Chen J, Luo Y, Guo H, Jiang RD, Liu MQ, Chen Y, Shen XR, Wang X, Zheng XS, Zhao K, Chen QJ, Deng F, Liu LL, Yan B, Zhan FX, Wang YY, Xiao GF, Shi ZL. 2020. A pneumonia outbreak associated with a new coronavirus of probable bat origin. Nature 579:270–273.

2. Wu F, Zhao S, Yu B, Chen YM, Wang W, Song ZG, Hu Y, Tao ZW, Tian JH, Pei YY, Yuan ML, Zhang YL, Dai FH, Liu Y, Wang QM, Zheng JJ, Xu L, Holmes EC, Zhang YZ. 2020. A new coronavirus associated with human respiratory disease in China. Nature 579:265–269.

3. WHO. 2020. World Health Organization - Coronavirus disease 2019 (COVID-19). Situation Report – 106. https://wwww.hoint/emergencies/diseases/novel-coronavirus-2019/situation-reports/ Accessed 21 Mar 2020.

4. Xie M, Chen Q. 2020. Insight into 2019 novel coronavirus - An updated interim review and lessons from SARS-CoV and MERS-CoV. Int J Infect Dis 94:119–124.

5. Tay MZ, Poh CM, Renia L, MacAry PA, Ng LFP. 2020. The trinity of COVID-19: immunity, inflammation and intervention. Nat Rev Immunol doi:10.1038/s41577-020-0311-8.

6. Sanders JM, Monogue ML, Jodlowski TZ, Cutrell JB. 2020. Pharmacologic Treatments for Coronavirus Disease 2019 (COVID-19): A Review. JAMA doi:10.1001/jama.2020.6019.

7. Elbein AD. 1991. Glycosidase inhibitors: inhibitors of N-linked oligosaccharide processing. FASEB J 5:3055–3063.

8. Fischl MA, Resnick L, Coombs R, Kremer AB, Pottage JC, Jr., Fass RJ, Fife KH, Powderly WG, Collier AC, Aspinall RL, et al. 1994. The safety and efficacy of combination N-butyl-deoxynojirimycin (SC-48334) and zidovudine in patients with HIV-1 infection and 200-500 CD4 cells/mm3. J Acquir Immune Defic Syndr (1988) 7:139–147.

9. Tierney M, Pottage J, Kessler H, Fischl M, Richman D, Merigan T, Powderly W, Smith S, Karim A, Sherman J, et al. 1995. The tolerability and pharmacokinetics of N-butyl-deoxynojirimycin in patients with advanced HIV disease (ACTG 100). The AIDS Clinical Trials Group (ACTG) of the National Institute of Allergy and Infectious Diseases. J Acquir Immune Defic Syndr Hum Retrovirol 10:549–553.

10. Dwek RA, Butters TD, Platt FM, Zitzmann N. 2002. Targeting glycosylation as a therapeutic approach. Nat Rev Drug Discov 1:65–75.

11. Chang J, Warren TK, Zhao X, Gill T, Guo F, Wang L, Comunale MA, Du Y, Alonzi DS, Yu W, Ye H, Liu F, Guo JT, Mehta A, Cuconati A, Butters TD, Bavari S, Xu X, Block TM. 2013. Small molecule inhibitors of ER alpha-glucosidases are active against multiple hemorrhagic fever viruses. Antiviral Res 98:432–440.

12. Wu SF, Lee CJ, Liao CL, Dwek RA, Zitzmann N, Lin YL. 2002. Antiviral effects of an iminosugar derivative on flavivirus infections. J Virol 76:3596–3604.

13. Platt FM, d’Azzo A, Davidson BL, Neufeld EF, Tifft CJ. 2018. Lysosomal storage diseases. Nat Rev Dis Primers 4:27.

14. Licastro D, Rajasekharan S, Dal Monego S, Segat L, D’Agaro P, Marcello A, Regione FVGLGoC. 2020. Isolation and full-length genome characterization of SARS-CoV-2 from COVID-19 cases in Northern Italy. J Virol doi:10.1128/JVI.00543-20.

15. Stadlbauer D, Amanat F, Chromikova V, Jiang K, Strohmeier S, Arunkumar GA, Tan J, Bhavsar D, Capuano C, Kirkpatrick E, Meade P, Brito RN, Teo C, McMahon M, Simon V, Krammer F. 2020. SARS-CoV-2 Seroconversion in Humans: A Detailed Protocol for a Serological Assay, Antigen Production, and Test Setup. Curr Protoc Microbiol 57:e100.

16. ter Meulen J, van den Brink EN, Poon LL, Marissen WE, Leung CS, Cox F, Cheung CY, Bakker AQ, Bogaards JA, van Deventer E, Preiser W, Doerr HW, Chow VT, de Kruif J, Peiris JS, Goudsmit J. 2006. Human monoclonal antibody combination against SARS coronavirus: synergy and coverage of escape mutants. PLoS Med 3:e237.

17. Tian X, Li C, Huang A, Xia S, Lu S, Shi Z, Lu L, Jiang S, Yang Z, Wu Y, Ying T. 2020. Potent binding of 2019 novel coronavirus spike protein by a SARS coronavirus-specific human monoclonal antibody. Emerg Microbes Infect 9:382–385.

18. Petris G, Bestagno M, Arnoldi F, Burrone OR. 2014. New tags for recombinant protein detection and O-glycosylation reporters. PLoS One 9:e96700.

19. Carletti T, Zakaria MK, Faoro V, Reale L, Kazungu Y, Licastro D, Marcello A. 2019. Viral priming of cell intrinsic innate antiviral signaling by the unfolded protein response. Nat Commun 10:3889.

20. Rajasekharan S, Marcello A. 2020. Multitarget synthetic RNA for SARS-Cov-2 detection and quantification. (unpublished).

21. Ritchie G, Harvey DJ, Feldmann F, Stroeher U, Feldmann H, Royle L, Dwek RA, Rudd PM. 2010. Identification of N-linked carbohydrates from severe acute respiratory syndrome (SARS) spike glycoprotein. Virology 399:257–269.

22. Fukushi M, Yoshinaka Y, Matsuoka Y, Hatakeyama S, Ishizaka Y, Kirikae T, Sasazuki T, Miyoshi-Akiyama T. 2012. Monitoring of S protein maturation in the endoplasmic reticulum by calnexin is important for the infectivity of severe acute respiratory syndrome coronavirus. J Virol 86:11745–11753.

23. Zakaria MK, Carletti T, Marcello A. 2018. Cellular Targets for the Treatment of Flavivirus Infections. Front Cell Infect Microbiol 8:398.

24. van Giersbergen PL, Dingemanse J. 2007. Influence of food intake on the pharmacokinetics of miglustat, an inhibitor of glucosylceramide synthase. J Clin Pharmacol 47:1277–1282.

25. Helenius A, Aebi M. 2001. Intracellular functions of N-linked glycans. Science 291:2364–2369.

